# Lifespan Normative Models of White Matter Fractional Anisotropy: Applications to Early Psychosis

**DOI:** 10.1101/2024.12.11.627897

**Authors:** Ramona Cirstian, Natalie J. Forde, Gary Zhang, Gerhard S. Hellemann, Christian F. Beckmann, Nina V. Kraguljac, Andre F. Marquand

## Abstract

This study presents large-scale normative models of white matter (WM) organization across the lifespan, using diffusion MRI data from over 25,000 healthy individuals aged 0-100 years. These models capture lifespan trajectories and inter-individual variation in fractional anisotropy (FA), a marker of white matter integrity. By addressing non-Gaussian data distributions, race, and site effects, the models offer reference baselines across diverse ages, ethnicities, and scanning conditions. We applied these FA models to the HCP Early Psychosis cohort and performed a multivariate analysis to map symptoms onto deviations from multimodal normative models using multi-view sparse canonical correlation analysis (msCCA). Our results reveal extensive white matter heterogeneity in psychosis, which is not captured by group-level analyses, with key regions identified, including the right uncinate fasciculus and thalami. These normative models offer valuable tools for individualized WM deviation identification, improving precision in psychiatric assessments. All models are publicly available for community use.

**Teaser:** Lifespan models of white matter offer insights into brain health, providing tools for tracking individual deviations across ages.

## Introduction

Over the past century, normative growth charts have become integral to paediatric practice, providing essential benchmarks for comparing individual growth patterns (height, weight, head circumference) with established population standards. These charts have facilitated a better understanding of typical developmental trajectories and have been crucial in identifying deviations from expected growth patterns which are used in clinical practice to determine if additional medical workup or treatment is required [1]. This concept has recently been extended to the field of neuroimaging, where it allows for detailed, individual-level insights into lifespan trajectories of brain measures. By comparing individual neuroimaging data against large, normative reference datasets, researchers and clinicians can gain a deeper understanding of both typical and atypical brain development and aging [2], [3], [4], [5].

In psychiatric disorders, traditional case-control studies have been valuable for detecting abnormalities in structural, microstructural, functional and neurometabolic brain signatures in patient groups compared to control groups. However, group comparisons are not designed to capture inter-individual heterogeneity which is prominent at the phenotypic and biological levels in virtually all psychiatric disorders. This significant translational gap hampers identification of specific biological markers that explain clinical heterogeneity in these disorders such as disease risk, severity, and progression, as well as responsiveness to pharmacological and non-pharmacological treatments and overall clinical outcomes. Normative modelling provides a precision framework that has emerged as a promising tool in this endeavour [6], [7], [8]. By comparing brain imaging data against large reference cohorts, this method allows us to quantify deviations from expected norms at the individual level. It is now possible to capture deviation profiles in a single patient, which offers a more nuanced understanding of biological variations in psychiatric disorders. Even more importantly, it also has promise for bridging this translational gap by providing a foundational framework for developing tailored tools that capture disease risk and progression, as well as precision treatments tailored to individual brain pathology. For instance, normative models capture inter-individual biological variations that provided important insights into heterogeneity in schizophrenia, major depressive disorder, bipolar disorder, ADHD and autism spectrum disorders [6], [9], [10]. Moreover, we have demonstrated that normative measures frequently outperform raw measures (e.g. cortical thickness in mm) in group difference testing, disease classification [11] and treatment response prediction [12].

We and others have created large-scale normative models that leveraged >50,000 healthy volunteer imaging datasets for structural [4], [5], [13] and functional MRI [11], [14]. To our knowledge, no comprehensive normative models for diffusion weighted imaging measures at comparable scale exist at this time. There are several reasons why this is the case. First, diffusion imaging was developed more recently than structural and functional MRI, and a broader adoption in neuroscience research did not happen until the early 2010s. Second, processing of diffusion data is more computationally demanding compared to structural and functional MRI and the gold standard for diffusion data quality control remains visual inspection; both of these factors have been substantial limitations to scaling efforts. Third, diffusion imaging measures are very sensitive to differences between vendors, individual scanners (e.g. signal intensity variations, eddy currents), and sequence acquisition parameters (e.g. b-values), making it difficult to integrate different datasets necessary to develop lifespan normative models. To date, only two preliminary studies have fit normative models to diffusion weighted data [15],[16]. In [15], the authors used approximately 1,300 single-shell DTI datasets collected at eight different sites using the same vendor to test performance of different statistical methods within the normative framework and in [16] the authors focus principally on generating reference curves for data harmonisation.

The aims of this study are to: (i) develop normative models of Fractional Anisotropy (FA), the most widely used diffusion metric in neuroimaging [17], across major white matter tracts using a large dataset of over 25,000 healthy individuals across a broad age range. By using high-quality diffusion MRI data from the UK Biobank and the Human Connectome Project, we seek to establish robust models that capture lifespan trajectories of white matter organization; (ii) investigate white matter FA in early psychosis, a prototypical psychiatric disorder that is known to be highly heterogeneous in disease severity and course, as well as clinical symptom expression and clinical outcomes. Using the HCP Early Psychosis (HCP-EP) dataset [18], we aim to map both group level differences and individual deviations from the normative model in order to better understand individual variability in white matter integrity; (iii) we aim to illustrate the value of normative models for multi-modal data fusion, by combining FA deviations with cortical thickness and subcortical brain volume deviations with the goal to identify multi-modal biological signatures and specific white matter pathways in psychosis associated with different psychosis symptom domains. Finally, (iv) we release all models freely to the community via our existing open-source software platforms [19].

## Results

### Normative modelling

First, we assembled high-quality multi-shell diffusion data from five cohorts having closely matched acquisition and processing pipelines, namely Human Connectome Project (HCP) Baby [20], HCP Development [21], HCP Young Adult [22], HCP Aging [23] datasets, and UK Biobank [24]. Total N=24,915, (N=12,457 for training and N=12,457 for test, stratified for sex, self-reported race, dataset and site). A summary of the sample and processing is provided in Figure 1 with further details in the methods. In short, the datasets were processed using harmonised FSL-based pipelines, involving pre-processing (intensity normalisation, distortion and movement corrections), DTI modelling to extract fractional anisotropy (FA) values, Tract-Based Spatial Statistics (TBSS) for skeletonised FA images, and segmentation with the Johns Hopkins University (JHU) atlas to compute mean FA values across 48 white matter tracts. We then fit lifespan normative models to these data on the basis of age, sex, site and race using warped Bayesian linear regression (BLR) and a non-linear basis expansion over age, in line with our prior work [4], [25]. We assessed the quality of the normative modeling fit using three key out-of-sample metrics, namely explained variance (EV), evaluating the fit of the median regression line, in addition to skewness and kurtosis, which evaluate the shape of the distribution used to model the centiles. These metrics offer insight into how well the models capture the underlying distribution of the data across 48 white matter tracts. The mean (standard deviation) EV was 0.37 (0.10), indicating good fit across different models. Skewness, and kurtosis were respectively -0.09 (0.12) and 0.42 (0.27), which together indicate that the shape was also appropriate for the data. Supplementary figure 1 shows a histogram of the EV, skew and kurtosis of the models.

**Figure 1.**
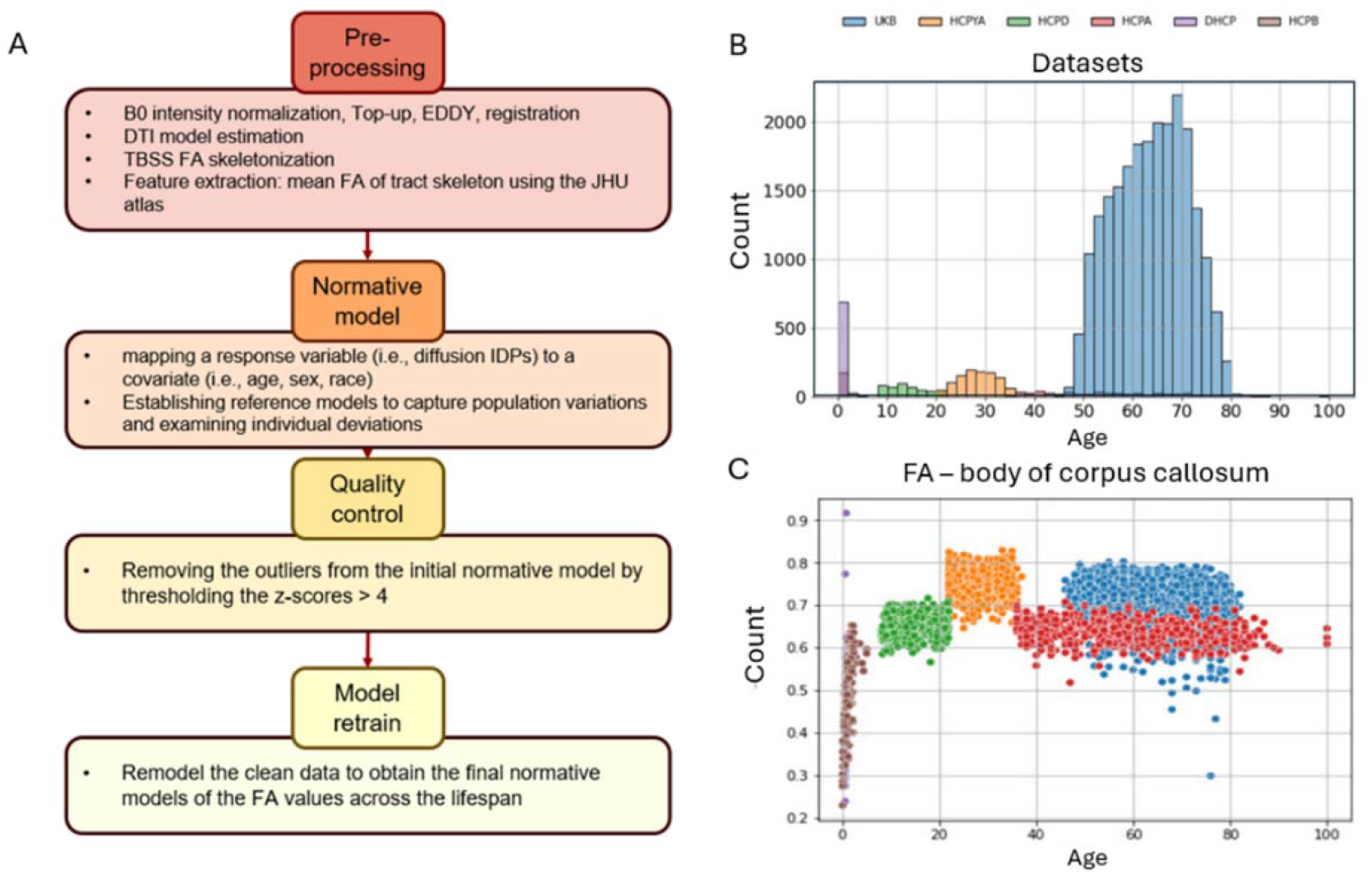
A) Flow chart of the main diffusion image processing steps B) Histogram plot of the data used for normative modeling, showing the population density at each age and highlighting the different datasets used C) Scatterplot exemplifying the quality control process using normative modeling and outlier exclusion based on Z-score thresholding. In this plot, site effects are clearly evident, which are accommodated by the normative models (see Figure 2). https://github.com/ramonacirstian/fa_normative_modelingn.d

We illustrate the trajectory and fitted centiles for a selection of white matter tracts across the lifespan in Figure 2. The complete set can be found in the supplementary figure 2. In addition, we also show the results of models that do not include race in the supplementary figure 3.

**Figure 2.**
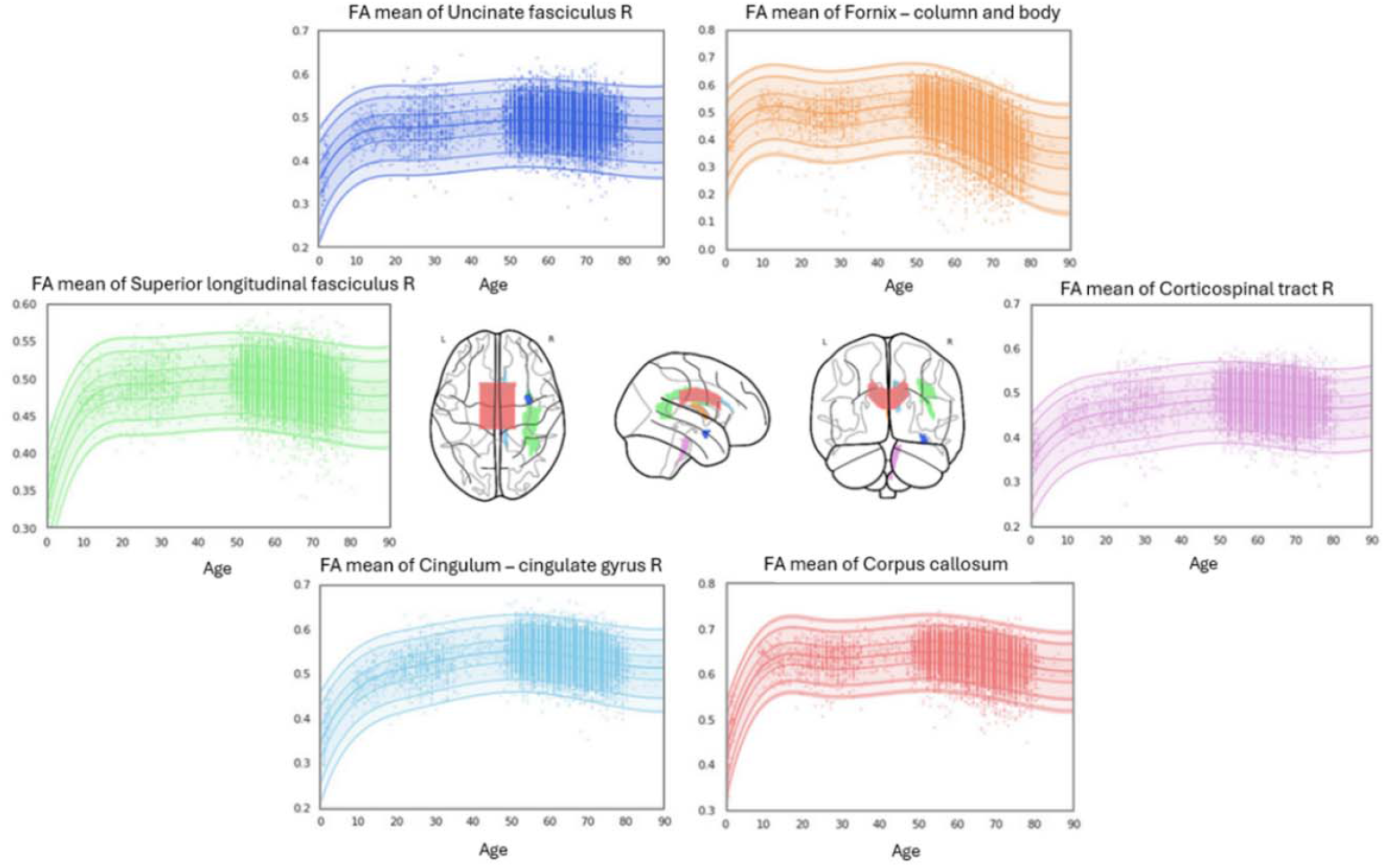
A selection of six white matter tracts and their corresponding normative modelling centile plots highlighting the similarity in white matter formation and degeneration along the lifespan as well as tract specific differences in terms of shapes and variance of the FA values. For visualization purposes, data from different sites are aligned to a common reference (e.g. the mean centiles or the centiles for an arbitrary chosen site) by computing the z-scores separately for each site using the site-specific means and standard deviations, then inverting the z-scores using the mean and standard deviation derived from the common reference.

### Application to a clinical dataset

Next, we used these models to understand heterogeneity in white matter FA in psychosis. To achieve this, we applied these reference models to the HCP early psychosis (HCP-EP) dataset (N=173 with diffusion data - see supplementary table 2 for demographic information) in order to derive z-scores for each individual and tract. We evaluated the mean differences in normative deviations between patients and controls for each tract using a t-test, applying false discovery rate (FDR) correction [26] to account for multiple comparisons. There were no significant differences in the mean deviations between individuals with psychosis and healthy controls that survived false discovery rate (FDR) multiple comparison correction, although we did find nominally significant effects in the fornix (column and body and the stria terminalis bilaterally). However, we did find evidence for significantly more heterogeneity in individuals with psychosis relative to controls in terms of the proportion of extreme deviations. More specifically, individuals with schizophrenia had a greater proportion of extreme positive (Mann-Whitney U=1403.0, p=0.0036) and extreme negative (U=1517.0, p=0.0016) Z-scores relative to controls, indicating substantial differences between groups that were highly variable across individuals. Notably, while both positive and negative deviations were present, the prominence of negative outliers (subjects with Z-scores exceeding ±2.6 was particularly pronounced, highlighting a consistent trend where patients exhibited a greater number of extreme Z-scores across white matter tracts. We show the percentage overlap of extreme deviations across all tracts in Figure 3, which reveals that extreme positive and negative deviations were observed in some individuals with psychosis in nearly all tracts. In contrast, the extreme deviations in controls were more focused, confined to only several white matter tracts, as illustrated in the bar plots in the supplementary figure 4, with an alternative representation highlighting overlap in individual tracts.

**Figure 3:**
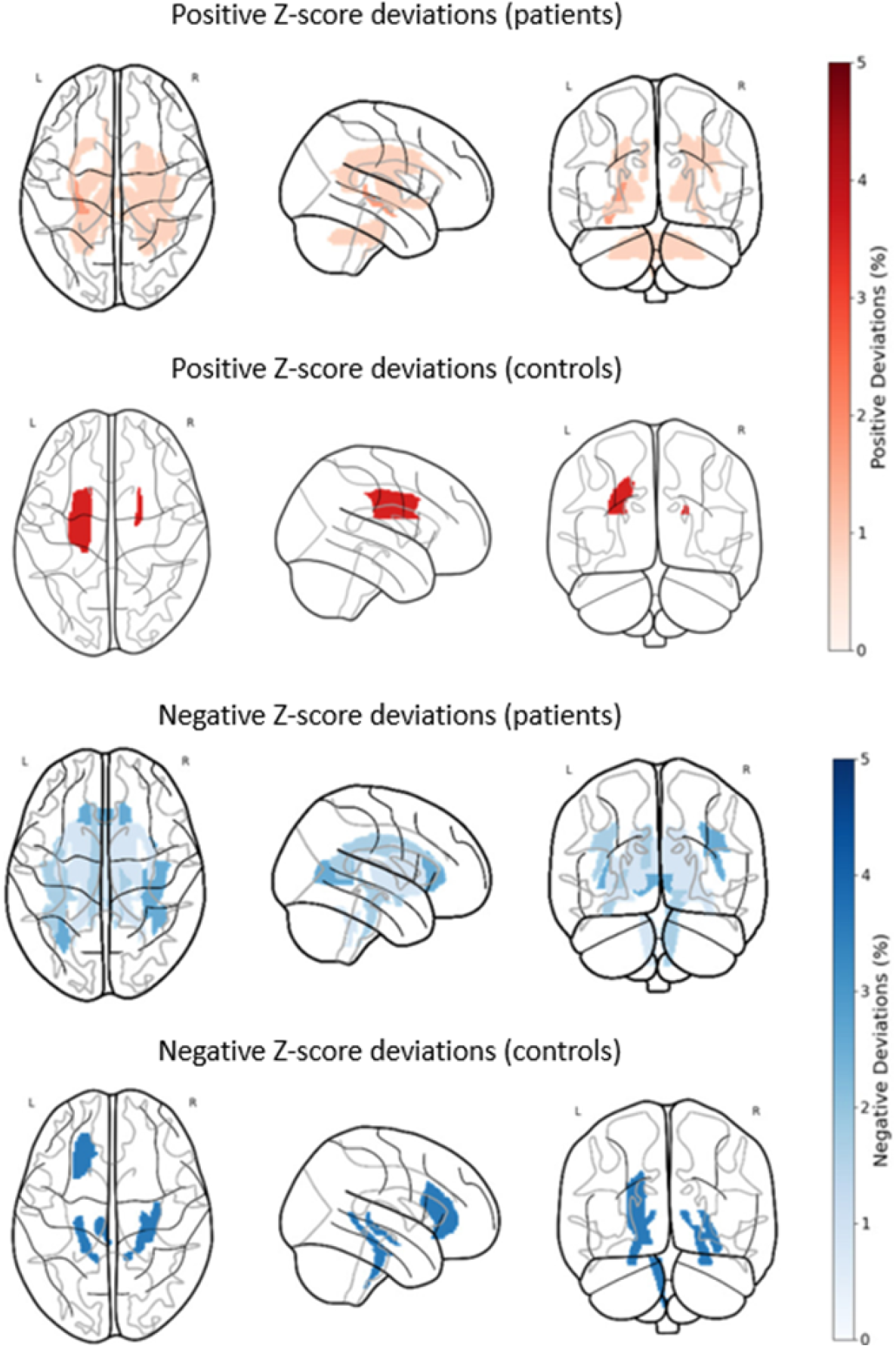
Glass brain representations illustrating the overlap between extreme positive and negative Z-score deviations for patients and controls, with thresholds set at - 2.6 and 2.6 which correspond to a p-value of 0.01. This stringent threshold enhances the detection of significant deviations while controlling for false positives. The top two panels depict positive Z-score deviations for patients and controls, while the bottom two panels show negative Z-score deviations for patients and controls. The legend indicates the percentage of subjects having extreme deviations in each tract.

Next, we sought to demonstrate the utility of these normative models in identifying multivariate brain-behaviour associations within a clinical cohort. To achieve this, we combined FA deviations with cortical thickness and subcortical volume measures from our previously published models [4] in a multimodal analysis. Symptom severity was quantified using the Positive and Negative Syndrome Scale (PANSS) [27], with domain scores for positive, negative, and cognitive symptoms, as well as the total score, summarised using a standard factor model, the ‘Marder’ factors, which were estimated and released by the HCP-EP consortium. More specifically, we included the positive symptom factor, negative symptom factor, cognitive/disorganised symptom factor in addition to the total PANSS score. To determine the multivariate association with symptoms, we used an approach we have employed in prior work [28], based on a multi-view sparse canonical correlation analysis (msCCA) and stability selection [29] (see methods for details). Briefly, aimed to learn the association between three ‘views’ of the data, namely symptom domains, FA deviations and structural deviations (i.e. deviations from normative models of cortical thickness and subcortical volume). Next, we randomly split the data 1000 times into training (70%) and test (30%) sets, then fit an msCCA model and report the mean canonical correlation on the test set. This analysis yielded a significant mean test canonical correlation of r=0.25 for the leading component (p=0.003 under permutation testing, see Methods for details). This model showed good predictive performance for both the associations between symptoms and FA deviations and symptoms and structural (cortical and subcortical) deviations, but not between diffusion and structural deviations (Figure 4 A). This is expected because we deliberately do not optimise directly for this to prevent the model learning the trivial correlation between different types of brain features (see methods). The second and third components achieved test canonical correlations of r=0.04 (p=0.11) and r=0.02 (p=0.03) respectively. However, considering the limited clinical relevance of associations of this magnitude and their marginal significance of the third component, we focus principally on the first.

**Figure 4:**
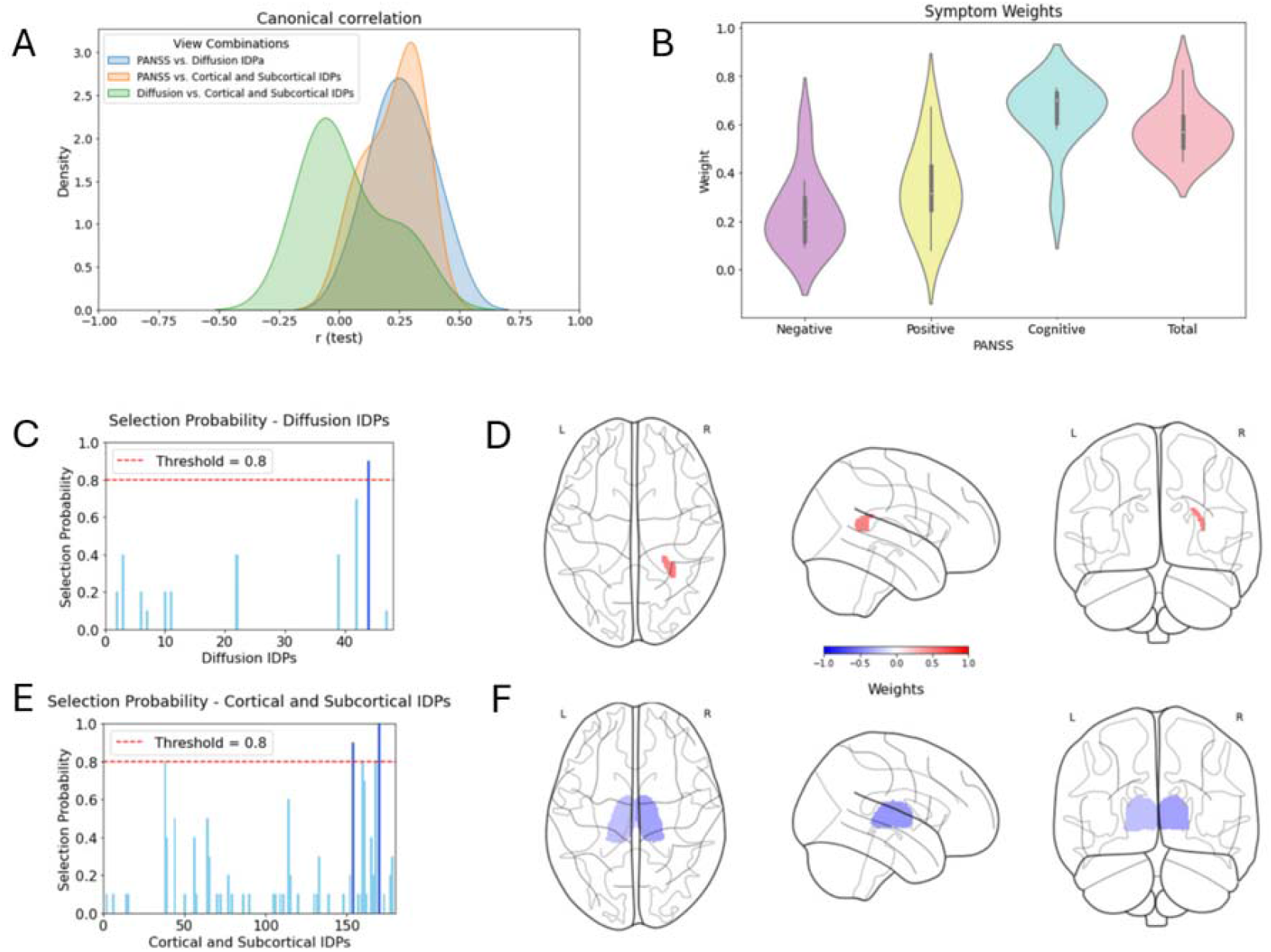
(A) Density plot of the multiple sparse Canonical Correlation Analysis (msCCA) main components, highlighting the distribution of test canonical correlations separately for each pair of views. (B) Violin plots representing the weights of PANSS symptom scores across the four symptom categories, namely negative symptoms, positive symptoms, cognitive symptoms, and total symptoms. (C) and (D) Selection probabilities for diffusion white matter tracts and Cortical Thickness white matter tracts, respectively, with a red threshold line indicating the chosen selection threshold of 80%. (E) and (F) Glass brain representations of the significantly selected white matter tracts and subcortical regions of interest, respectively. Note that no cortical thickness ROIs survived the selection threshold. The highlighted regions include the uncinate fasciculus (right) for diffusion and the cortical thickness regions: Left-Thalamus and Right-Thalamus

The symptom loadings derived from the msCCA analysis show that the association was principally driven by the cognitive factor and total PANSS scores (Figure 4 B). We used stability selection to determine the most informative features driving the association by counting the number of times each feature was selected under the 1000 random splits described above and considered samples having a selection probability greater than 0.8 as informative. Note that this threshold is theoretically justified in order to control the type 1 error rate [29]. Under this threshold, FA in the right uncinate fasciculus and volume of the thalamus bilaterally were predictive of PANSS symptoms Figure 4 C-F.

## Discussion

This study presents a set of large-scale normative models for FA across major white matter tracts, estimated from a dataset of over 25,000 individuals spanning infancy to old age. Leveraging high-quality multi-shell diffusion MRI data, these models map the trajectory of white matter development and degeneration over the lifespan whilst also quantifying variance across the population. We showcase the clinical utility of these models by mapping inter-individual variation in cohorts of individuals with early psychosis. We show a high degree of inter-individual heterogeneity in these individuals, evidenced by relative increases in both positive- and negative deviations from the normative model in individuals with psychosis relative to controls. These differences were evident despite an absence of case control effects, indicating that the differences were highly individualized. Finally, we show that normative deviations of FA, cortical thickness and subcortical brain volume were accurate multi-modal predictors of symptomatology. Taken together, our findings provide a step toward advancing the understanding of the heterogeneity of white matter alterations in early psychosis.

Our normative models show region-specific developmental trajectories in white matter organization that align well with foundational findings on lifespan changes in FA [30], [31]. However, we also show that inter-individual variability is considerably higher than the magnitude of lifespan-related changes in FA, underscoring the importance of using approaches such as this to characterize this at the individual level. Studies suggest that increased FA during development relates to synaptic pruning and myelination, while declines in old age are linked to axonal degradation and reduced fiber coherence [32], [33]. Our models robustly capture these patterns, underscoring their relevance as a normative reference sample and utility for studies examining brain aging and clinical conditions.

In the HCP-EP cohort, we show a high degree of inter-individual variability in white matter organization in psychosis, consistent with the variability that has already been described in brain structure [9], [10], and in other psychiatric conditions [6], [7], [34], which speaks to the potential for normative models as a basis for stratifying cohorts [3]. Note that this variability was evident in an absence of case-control effects, which indicates that inter-individual variability masks group level effects, which we also have observed in gray matter in autism [7]. We also show a multivariate correspondence between brain connectivity deviations, structural deviations and clinical symptoms driven by decreased volume in the thalamus and FA in the right uncinate fasciculus. The left and right thalamus showed negative weights, indicating that reductions in subcortical volume may be linked to greater symptom severity. The uncinate fasciculus exhibited a positive weight in relation to clinical symptoms, suggesting a possible compensatory role, although we cannot rule out that this finding may also reflect other factors (such as crossing fibres). In line with this interpretation, alterations in the uncinate fasciculus have been previously reported in psychotic disorders, suggesting its involvement in the pathophysiology of these conditions [14], [35]. Studies utilizing DTI have reported abnormalities in the uncinate fasciculus among individuals with schizophrenia and affective psychosis. For instance, Kawashima et al. [36] found reduced FA in the uncinate fasciculus of patients with recent-onset schizophrenia, indicating compromised white matter organisation in this tract [36].

One of the benefits of this study is that we focus on acquiring a high-quality diffusion sample with closely harmonized protocols. This maximizes the ability to attribute detected variations to biological differences, rather than artefacts such as data quality or residual site effects. In this study, we prioritised modelling FA as it is the most commonly used diffusion metric in the field due to its sensitivity to microstructural integrity factors like axonal density, fibre coherence, and myelination, making it a valuable and accessible measure for understanding white matter architecture. Additionally, FA is less affected by CSF contamination compared to metrics like mean diffusivity (MD), allowing for more accurate assessments in regions prone to such contamination, such as the fornix [37]. However, this is only the first step, we intend to augment these models with further models, including other tensor-based metrics, such as mean diffusivity MD and non-tensor models (e.g. neurite orientation dispersion and density imaging; NODDI [38], [39], to take full advantage of the multi-shell diffusion data and provide an even more comprehensive resource for white matter analysis. Finally, we provide these models to the field via our established no-code software platform [19] and via open-source software tools (https://github.com/ramonacirstian/fa_normative_modeling), so that others in the field can easily apply these models to their own data.

Finally, we acknowledge some limitations to the current study. The age distribution in our dataset is skewed, with fewer data points at the extremes of the lifespan particularly in young children between the ages of 5 and 8 and adults over 85 years old. Although great care was taken to ensure that this did not bias the analysis (e.g. by ensuring smoothness for the interpolating centile curves), this gap should be considered as limiting the generalizability of the models for younger and older populations at this time. We intend to augment our dataset with additional samples to increase data density in these regions as future high-quality datasets come online. Additionally, while our models effectively account for site-specific differences, variability due to demographic factors like socioeconomic background was not fully explored and should be considered in future normative modeling efforts. A strength of our analysis is that we specifically account for ethnicity in our models, by including self-reported race using fixed effects in the analysis, following our prior work [40]. We consider this important to reduce the risk of racial bias, but it should be remembered that the datasets on which these models were trained on are not representative of the wider population and are themselves biased towards ‘Western Educated, Industrialised, Rich and Democratic’ (WEIRD) populations [40]. Self-reported race is also an imperfect proxy for ethnicity, and it is likely that using more flexible modelling approaches may be needed to properly account for these effects [41]. For these reasons we also release the models that do not include race so that each researcher using these models can decide for themselves which model is more appropriate for their needs.

In summary, this study provides comprehensive normative reference models for FA across the lifespan, using an extensive dataset that spans infancy to old age. By integrating high-quality diffusion MRI data and using robust modeling techniques, we captured the typical trajectory of white matter development and decline, aligning with prior research and enhancing the field’s understanding of brain aging. Our application of these models to a clinical early psychosis cohort underscores their potential utility in identifying atypical white matter patterns in psychiatric conditions. These models not only serve as a benchmark for individual-level assessments but also offer valuable insights for precision medicine, facilitating more personalized interventions. This study highlights the relevance of normative modeling in neuroimaging, paving the way for its integration into clinical and research settings focused on individual variability in brain structure and pathology.

## Materials and Methods

### Data acquisition and processing

The construction of the lifespan dataset involved integrating data from five cohorts having high-quality multi-shell diffusion data, i.e.: the HCP Baby [20], HCP Development [21], HCP Young Adult [22], HCP Aging [23] datasets, and the UK Biobank [24]. The demographic information is available in supplementary table 1.

The processing of these datasets followed harmonized FSL-based pipelines, summarized in Figure 1A. Initially, pre-processing was performed: B0 intensity normalization, correction for EPI distortions, eddy-current-induced and movement corrections. These corrections were executed using the HCP-pipeline [42] for the HCP datasets while the UKB dataset was already processed according to the UKB documentation [43]. Subsequently, we estimated the DTI model using DTIfit on the lowest shell value in order to extract the fractional anisotropy (FA) values. Following this, we ran Tract-Based

Spatial Statistics (TBSS) [44] on the FA images which included registration to a standard space (FMRIB58_FA), projection of each individual’s FA image to the standard space skeletonized image (threshold at 0.2) to generate skeletonized FA images for each individual in the same space. Finally, segmentation was conducted using the Johns Hopkins University (JHU) atlas [45]. This process delineated 48 white matter (WM) tracts (listed the supplementary figure 4), for which we computed the mean FA values along the skeleton of each tract.

### Normative modeling

To prepare for the modelling stage, we began by splitting the dataset of subjects (N=24,915) into two equal groups: a test set (N=12,457) and a training set (N=12,457), stratified to ensure an even distribution of sex, race, dataset and site. A normative model was then fit to the training set for each white matter tract. The model incorporated several covariates, including sex, age, and dummy coded race, and site. To address potential non-linear effects and non-Gaussian distributions, we employed a warped Bayesian linear regression (BLR) model and used in previous research [4], [25]. This approach involved applying a third-order polynomial B-spline basis expansion over age, with five evenly spaced knots, combined with a SinhArcsinh warping function.

Next, we estimated deviation scores for each subject and white matter tract. In line with our prior work [46] we refit the models after excluding gross outliers having deviations larger than 5 standard deviations from the mean (Figure 1C). Once the models were refit with the cleaned data, we calculated the fit statistics, including explained variance, skew, and kurtosis. The extent of deviation for each subject was visualized by plotting individual z-scores against the mean and centiles of variation predicted by the model. All statistical analyses were conducted using Python version 3.8, with the Predictive Clinical Neuroscience PCN toolkit (GitHub, PCNtoolkit).

### Application to a clinical dataset

Next, we applied the model to the Human Connectome Project Early Psychosis (HCP-EP) dataset [18] (Jacobs et al., 2024), which includes multi-shell diffusion data and T1-weighted structural MRI derived from participants diagnosed with early psychosis (n=118) and control participants (n=55). The dataset’s demographic distribution comprises 37% females and 63% males, with a racial composition of 58% White, 28% Black, 9% Asian, 1% Mixed, and 3% Other. Participants with early psychosis were diagnosed using the Structured Clinical Interview for DSM-5 (SCID-5) (First et al., 2015) and symptoms assessed with the Positive and Negative Syndrome Scale (PANSS) [27], including negative symptoms (e.g., social withdrawal), positive symptoms (e.g., hallucinations), disorganisation, and general psychopathology. The item-level data were subsequently summarized by the HCP-EP consortium using a standard factor model [47] and the positive, negative and cognitive symptom domain scores were used in in addition to the PANSS total score to quantify symptomatology across multiple domains [47]. Medication status was also documented, including antipsychotic type and dosage converted to chlorpromazine equivalents.

The diffusion data were processed with the same pipeline as described above (Figure 1A), and structural data were processed using Freesurfer version 6.0 following similar procedures as we have described previously [4]. Next, we divided this dataset into a training set, consisting of half of the control participants, and combined it with the larger training set described above to retrain the normative models for each white matter tract. Using transfer learning, as in our previous work, we can efficiently adapt the models with only a small amount of calibration data to account for site-specific effect. We then computed z-scores for the patients and remaining controls for the FA data and computed the deviations for cortical thickness and subcortical volumes derived from models we have previously brought online [4]. Note that the splits for this analysis were matched so that the same participants were in the training and test sets for diffusion and structural measures at each iteration.

We next assessed the mean difference of the deviations between patients and controls for each tract using a t-test with false discovery rate (FDR) correction for multiple testing [26]. We then tested whether the proportion of extreme deviations differ between groups for each tract. To achieve this, we calculated the percentage of participants falling below and above the threshold in each of the 48 tracts. To achieve this, we set a z-score threshold between -2.6 and 2.6, which correspond to a p-=value of 0.01 as in prior work to identify extreme deviations then employed a non-parametric Mann-Whitney U test [48], again followed by FDR correction for multiple comparisons. This stringent threshold enhances the detection of significant deviations while controlling for false positives

Next, we combined the FA deviations with structural deviations in a multimodal analysis aiming to predict the four symptom domains described above. To achieve this, we conducted a multi-view sparse canonical correlation analysis (msCCA), using an approach we have described previously [28]. To identify relationships between multiple datasets, msCCA maximises the cross-correlation between weighted sums of variables from each dataset (Equation 1).

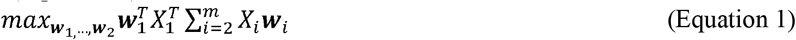

where *X*□ represents psychiatric symptoms and *X*□ to *X*_*m*_ represent neuroimaging measures from *m* different modalities. The weights (***w***□, ***w***□, …, ***w***_*m*_) are subject to constraints: |*W*_1_|_2_ = 1, |*W*_*i*_|_2_ = 1, |*W*_1_|_1_ *c*_1_, |*W*_*i*_ |_1_ *c*_*i*_, ensuring sparsity and interpretability. The regularization parameters for each view *v* are assumed to be set such that 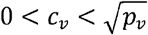 where *p*_*v*_ is the number of feature in view *v*, which is helpful to bound the (approximate) number of selected variables [28]. Importantly, this approach avoids optimising correlations between different neuroimaging modalities directly, focusing instead on shared variance between psychiatric symptoms and neuroimaging measures.

This involved creating three views of the data (i.e. symptoms, structural deviations and diffusion deviations) and then fitting an msCCA model to maximise the association between symptom domains and each of the diffusion and structural deviations but – crucially– not the imaging views with one another [28]. This requires setting (L1-norm) sparsity parameters for each of the data views *c*_*v*_. These were fixed throughout such that approximately 90% of the PANSS features were selected and 20% of the FA and structural image features. This corresponds respectively to light regularization for the symptoms, and moderately high regularization for the FA and structural measures. Note that we deliberately choose fixed parameters rather than optimizing them via nested cross-validation given the moderate sample size for the clinical data. Instead, we employed stability selection to assess the generalizability of the coefficients, which is theoretically guaranteed to provide tight type-I family-wise error control [29].

In more detail, we performed 1000 random splits of the dataset into a training (70%) and test set (30%) and selected the most stable features, i.e. features that were selected in more than 80% of the splits. This threshold is justified as it is sufficiently high that the theoretical guarantees on controlling the type 1 error rate become operative. In order to assess generalizability, we then ran an additional 1000 permutations, where within each permutation, we computed the test canonical correlation averaged across 10 random splits of the data, both before and after randomly permuting the order of the PANSS data view to destroy the relationship between the symptom scores and imaging data. We did this for the first three canonical components, which were derived by successively applying projection deflation to the data matrices [28], [49]. In order to compute significance, we then counted the number of times the true mean test canonical correlation exceeded the permuted value and divided by the number of permutations.

## Supporting information

supplemetary figure 1: normative model fit statistics

supplemetary figure 2: normative model centiles with race

supplemetary figure 3: normative model centiles without race

supplemetary figure 4: extreme deviations patients vs controls

supplemetary table 1: demographic information of all datasets

supplemetary table 2: demographic information of HCPEP

## Acknowledgments

NIH grant number 1R01MH130362-01A1

## Author contributions

Conceptualization: RC, NF, AM

Methodology: RC, NF, AM

Validation: RC, NF, AM, NK, GH

Software: RC, NF, AM

Formal analysis: RC, NF, AM

Visualization: RC, NF, AM

Supervision: NF, AM, CB

Writing—original draft: RC

Writing—review & editing: RC, NF, AM, NK, GZ

## Data and materials availability

The data used in the present study is part of the UK Biobank dataset which is available to be downloaded upon completing an access application. More information can be found on the dedicated webpage (UK Biobank, n.d.). The code used to process the data and train the normative models is also available online on GitHub (https://github.com/ramonacirstian/fa_normative_modelingn.d.)

