## Supplementary material for "Lifespan Normative Models of White Matter Fractional Anisotropy: Applications to Early Psychosis": supplemetary figure 1: normative model fit statistics

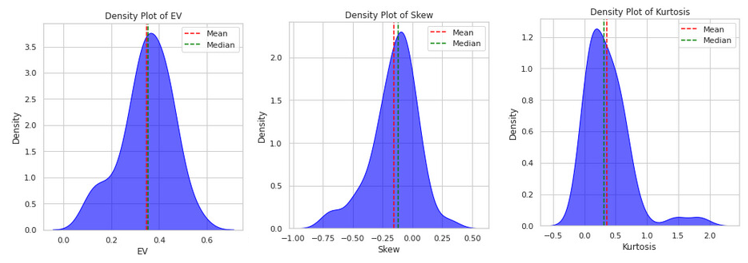
Fig. S1.

Supplementary figure 1: Density plots of the Explained Variance (EV), Skewness, and Kurtosis for the normative models fitted to 48 white matter tracts (FA) in the lifespan dataset. The red dashed lines indicate the mean values, and the green dashed lines represent the median values for each metric. These plots highlight the distributional properties of the normative models, providing insights into their variability and performance.
