## Supplementary material for "Lifespan Normative Models of White Matter Fractional Anisotropy: Applications to Early Psychosis": supplemetary figure 3: normative model centiles without race

Fig. S3.

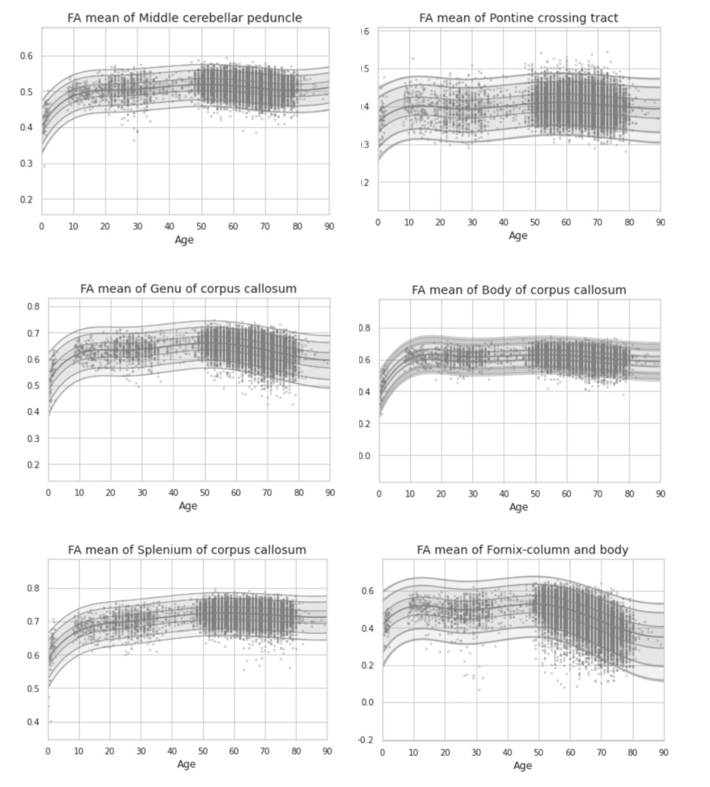

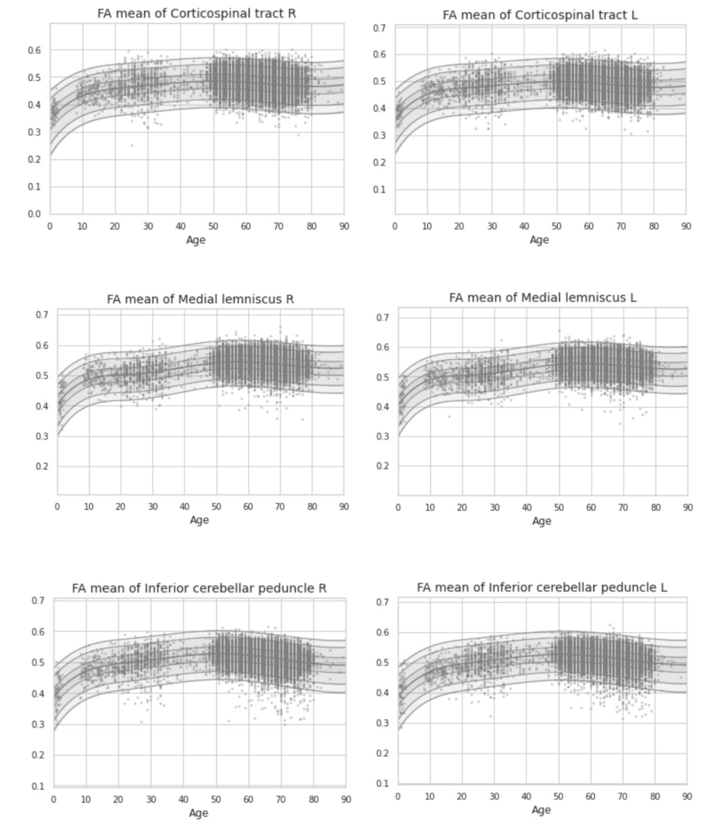

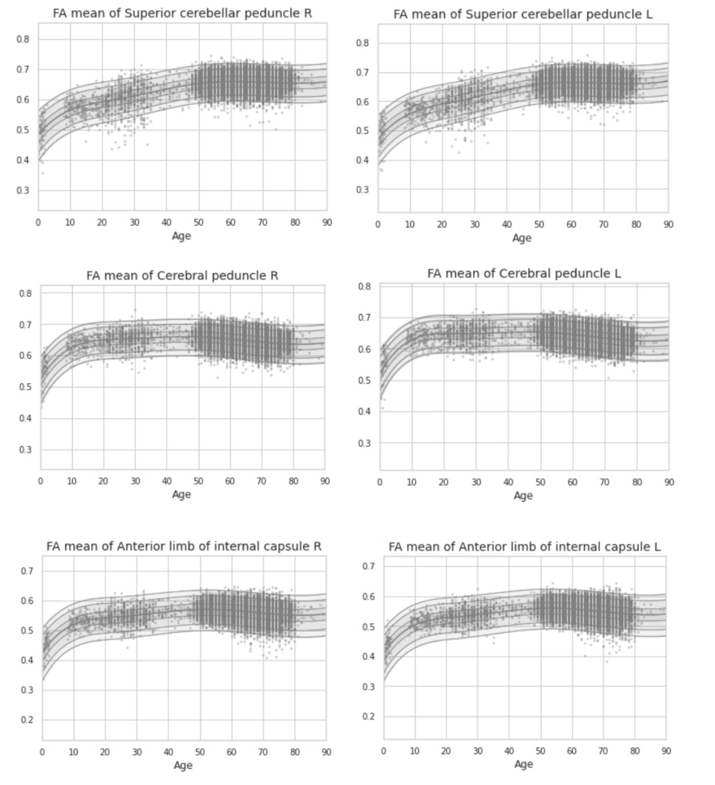

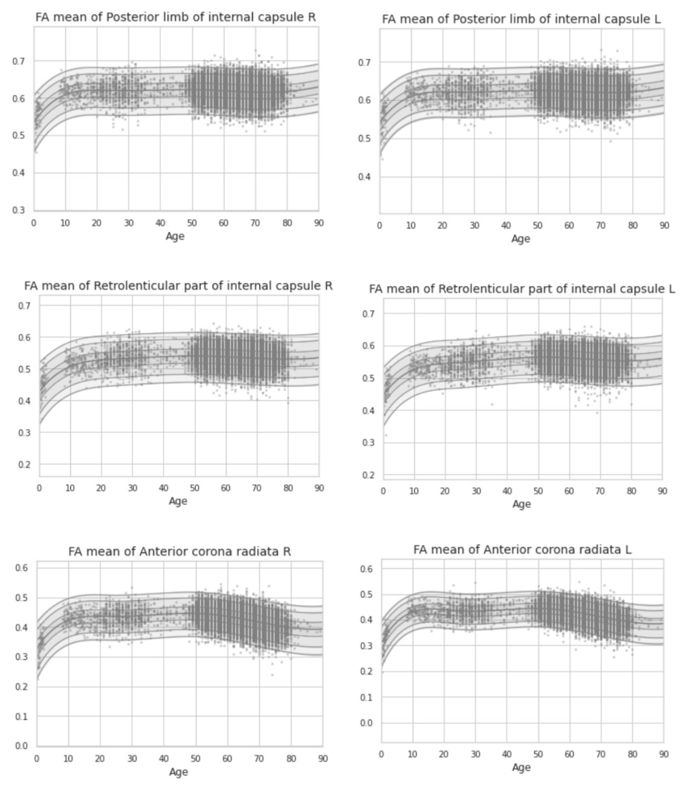

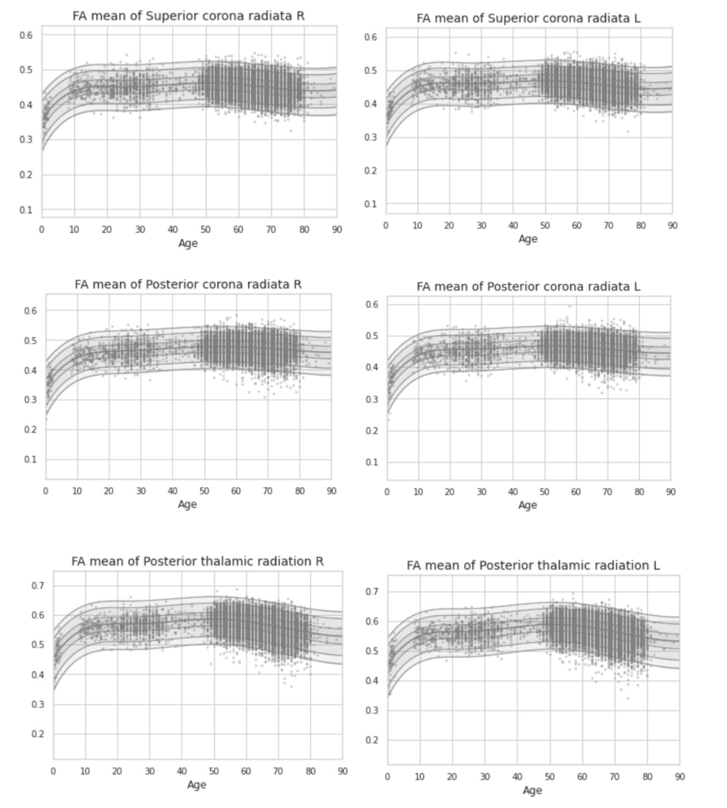

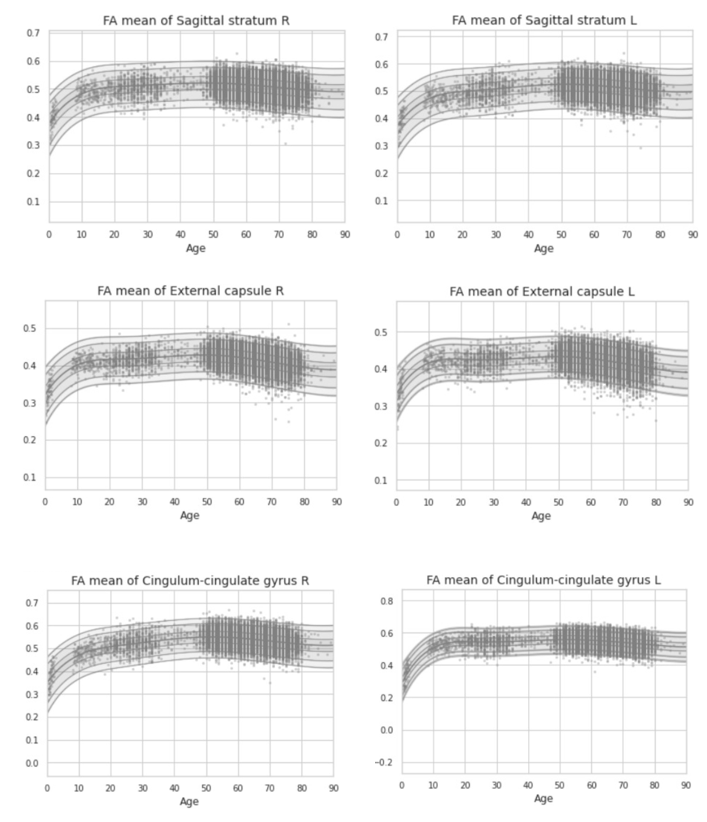

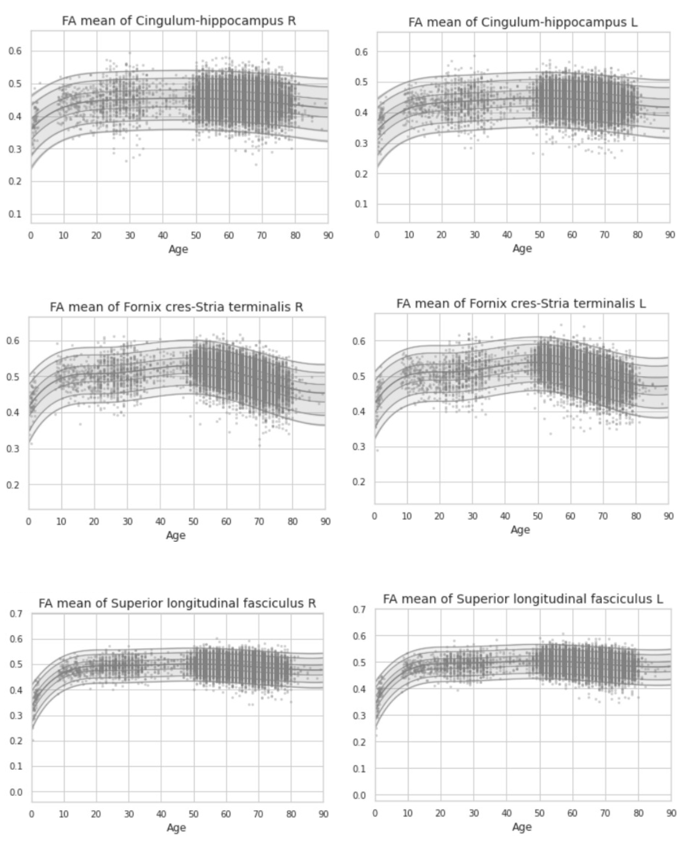

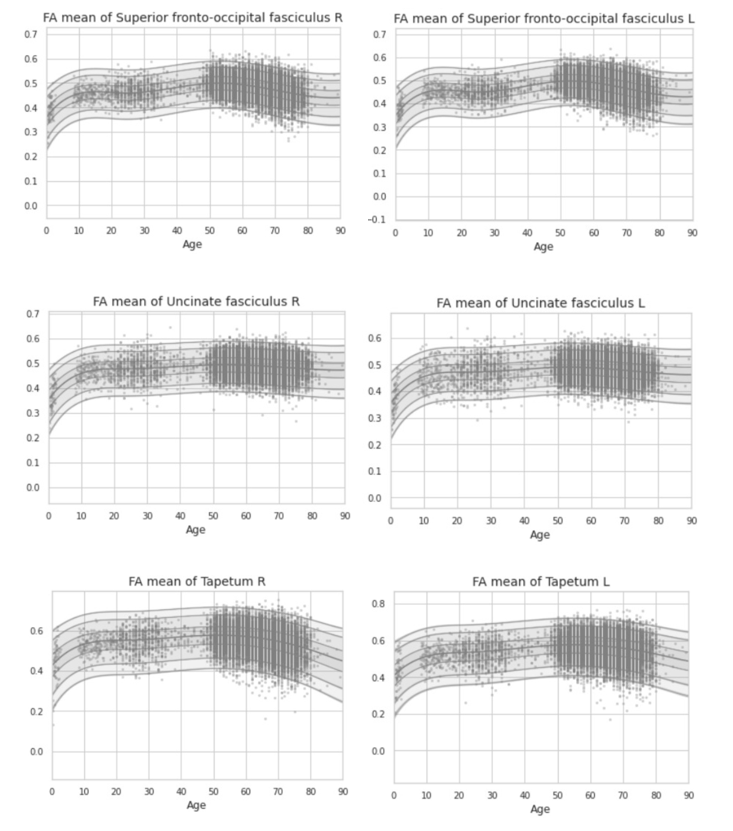

Supplementary figure 3: Centile plots illustrating the normative models for Fractional Anisotropy (FA) across 48 white matter tracts in the lifespan dataset. Each plot visualises the distribution of individual data points (grey dots) and the corresponding normative centiles (shaded bands), which capture the variability within the population. These models did not include race as a covariate. The centiles highlight age-related changes and variability in FA for each tract, offering insights into the developmental and degenerative patterns observed across the lifespan.
