## Supplementary material for "Lifespan Normative Models of White Matter Fractional Anisotropy: Applications to Early Psychosis": supplemetary figure 4: extreme deviations patients vs controls

Fig. S4.

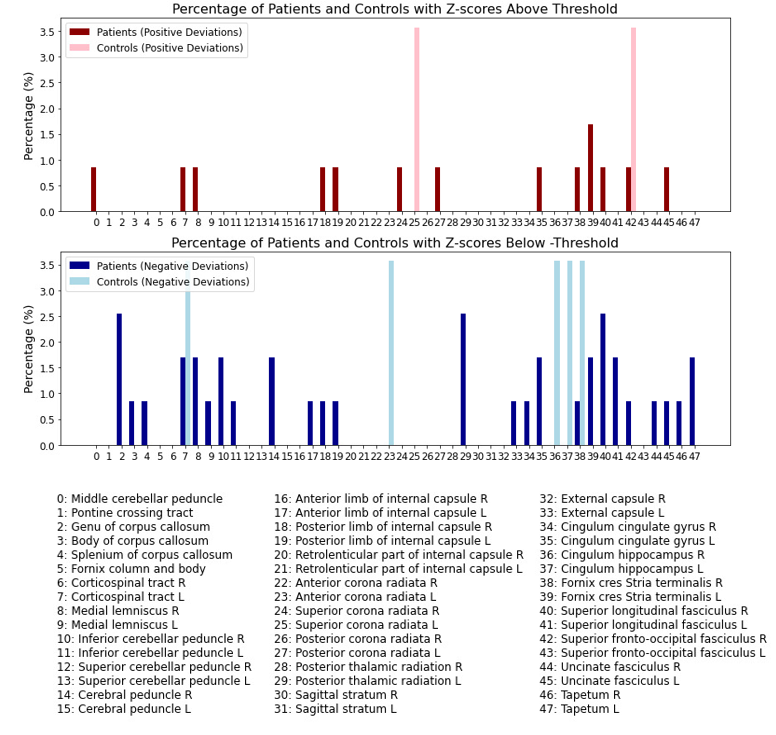

Supplementary figure 4: Bar plots showing the percentage of patients and controls identified as outliers with Z-scores greater than ±2.6, based on normative models fitted to 48 white matter tracts. The top plot represents positive deviations (Z-scores > 2.6) for patients and controls, while the bottom plot represents negative deviations (Z-scores < -2.6). The percentage of outliers is displayed for each white matter tract, listed in the legend below the plots. This figure highlights the variability in the proportion of extreme deviations across different tracts and between patient and control groups
