## Supplementary material for "Lifespan Normative Models of White Matter Fractional Anisotropy: Applications to Early Psychosis": supplemetary table 1: demographic information of all datasets

**Table S1**.

| **Dataset** | **Age (years)** | **Female** | **Male** | **Asian** | **Black** | **Mixed** | **Other** | **White** |
| --- | --- | --- | --- | --- | --- | --- | --- | --- |
| HCPA | 36 to 100 | 57.02% | 42.98% | 7.96% | 14.95% | 5.05% | 2.14% | 69.90% |
| HCPB | 0.04 to 3 | 53.37% | 46.63% | 2.81% | 3.37% | 11.80% | 18.54% | 63.48% |
| HCPD | 8 to 22 | 53.52% | 46.48% | 6.83% | 12.11% | 14.98% | 1.76% | 64.32% |
| HCPYA | 22 to 37 | 53.99% | 46.01% | 6.10% | 14.08% | 2.54% | 1.88% | 75.40% |
| UKB | 46 to 82 | 53.17% | 46.83% | 1.32% | 0.60% | 0.48% | 0.70% | 96.90% |

Supplementary table 1: Demographic characteristics of the datasets used to train and test the normative models. The table summarises the age ranges, gender distribution (percentage of female and male participants), and racial/ethnic composition (percentage of participants identifying as Asian, Black, Mixed, Other, and White) for each dataset: HCPA, HCPB, HCPD, HCPYA, and UKB. These datasets collectively span a wide range of ages and demographic diversity, ensuring robust model generalisability across populations. While all datasets are predominantly composed of white participants, some include a reasonable representation of Black, mixed-race, and Asian participants. The 'other' category encompasses individuals who either did not disclose their race or belong to a group too small to be categorized separately (e.g., American Indian/Alaska Native).
