## Supplementary material for "Lifespan Normative Models of White Matter Fractional Anisotropy: Applications to Early Psychosis": supplemetary table 2: demographic information of HCPEP

Table S2.

|  | Patients | Controls |
| --- | --- | --- |
| N | 118 | 28 |
| Age (μ, σ) | 22.7, 3.7 | 24.4, 4.0 |
| Sex (M%, F%) | 62%, 38% | 61%, 39% |
| Race (W%, B%, A%, Mx%, O%) | 51.7%, 38.1%, 5.9%, 0.85%, 3.4% | 75.0%, 7.1%, 14.3%, 0.0%, 3.6% |
| PANSS total score (μ, σ) | 47.0, 15.9 | NA |
| PANSS positive score (μ, σ) | 1.6, 0.9 | NA |
| PANSS negative score (μ, σ) | 2.0, 1.1 | NA |
| PANSS general score (μ, σ) | 1.5, 0.8 | NA |
| Marder positive score (μ, σ) | 12.5, 5.8 | NA |
| Marder negative score (μ, σ) | 14.0, 4.9 | NA |
| Marder cognitive score (μ, σ) | 10.7, 3.1 | NA |

Supplementary table 2. Demographic and Symptom Characteristics of the HCP Early Psychosis (EP) Dataset. This table summarises the demographic and clinical information for patients and controls in the HCP EP dataset. The total number of participants (N) is shown, along with the mean (μ) and standard deviation (σ) of their ages. Sex distribution is presented as the percentage of males (M%) and females (F%). Race is described by the percentage of participants identifying as White (W%), Black (B%), Asian (A%), Mixed (Mx%), or Other (O%). Symptom measures include the total PANSS score, positive, negative, and general PANSS scores, and Marder factor scores for positive, negative, and cognitive symptoms. For control participants, PANSS and Marder scores are not applicable (NA).
